# Manifold Learning Enables Interpretable Analysis of Raman Spectra from Extracellular Vesicle and Other Mixtures

**DOI:** 10.1101/2023.03.20.533481

**Authors:** Mohammadrahim Kazemzadeh, Miguel Martinez-Calderon, Robert Otupiri, Anastasiia Artuyants, Moi M. Lowe, Xia Ning, Eduardo Reategui, Zachary D. Schultz, Weiliang Xu, Cherie Blenkiron, Lawrence W. Chamley, Neil G.R. Broderick, Colin L. Hisey

## Abstract

Extracellular vesicles (EVs) have emerged as promising diagnostic and therapeutic candidates in many biomedical applications. However, EV research continues to rely heavily on in vitro cell cultures for EV production, where the exogenous EVs present in fetal bovine (FBS) or other required serum supplementation can be difficult to remove entirely. Despite this and other potential applications involving EV mixtures, there are currently no rapid, robust, inexpensive, and label-free methods for determining the relative concentrations of different EV subpopulations within a sample. In this study, we demonstrate that surface-enhanced Raman spectroscopy (SERS) can biochemically fingerprint fetal bovine serum-derived and bioreactor-produced EVs, and after applying a novel manifold learning technique to the acquired spectra, enables the quantitative detection of the relative amounts of different EV populations within an unknown sample. We first developed this method using known ratios of Rhodamine B to Rhodamine 6G, then using known ratios of FBS EVs to breast cancer EVs from a bioreactor culture. In addition to quantifying EV mixtures, the proposed deep learning architecture provides some knowledge discovery capabilities which we demonstrate by applying it to dynamic Raman spectra of a chemical milling process. This label-free characterization and analytical approach should translate well to other EV SERS applications, such as monitoring the integrity of semipermeable membranes within EV bioreactors, ensuring the quality or potency of diagnostic or therapeutic EVs, determining relative amounts of EVs produced in complex co-culture systems, as well as many Raman spectroscopy applications.

---

Extracellular vesicle (EV) research has grown rapidly in recent years due to the potential value that EVs could provide in diverse biomedical applications. Their protective lipid membranes, complex molecular cargo, and presence in all body fluids have encouraged development of EVs as liquid biopsy biomarkers or even as therapeutics. However, because patient samples are often too heterogeneous, limited, or simply unavailable, researchers have relied extensively on the use of in vitro cell cultures to produce a limitless supply of relatively homogeneous EVs. While the development of EV production and isolation techniques from cell cultures has played a crucial role in providing accessibility and advancing EV research worldwide, it has also revealed several issues. Among these is a strong dependence on serum supplements to support the growth of many types of cell cultures, leading to the eventual co-isolation of EVs of interest with exogenous EVs and particles (EVPs) [1, 2].

Several researchers have recently shown that exogenous EVs from serum supplements can have significant effects on experimental outcomes[3–9]. These findings have encouraged the development of mitigation strategies such as the use of commercially available EV-depleted serum, in-house EV depletion using centrifugation or filtration, serum starvation, and bioreactors that allow the use of serum supplements while excluding undesired EVs via semipermeable membranes[10–12]. However, few techniques can easily quantify or validate the purity of these EV preparations, with many researchers relying on simple nanoparticle counts or serum EV controls that may not adequately reflect the constituent EVs post-culture or post-isolation. Clearly, to improve the rigor and reproducibility in this growing field, a rapid, label-free, and transparent method of assessing the relative amounts of serum supplement-derived EVs within EV preparations from cell cultures would be ideal. While a few recent studies have made progress in using Raman spectroscopy-based methods to quantify the purity of EVs or determine the relative ratios of different EV or lipoprotein subpopulations, no system has presented a robust and sufficiently advanced analytical method which enables explainable interpretation[13–16].

Raman spectroscopy, particularly surface-enhanced Raman spectroscopy (SERS), may meet this need as it has enabled increasingly advanced applications in the chemical [17–21] and biological sciences [22–30] due to ongoing improvements in both instrumentation and analysis. The unique ability of SERS to non-destructively fingerprint chemical and molecular species within diverse samples makes it possible to carefully monitor changes in biological systems or other dynamic processes, provided that sufficiently advanced analytical tools can be applied to the acquired spectra[28–31]. In EV research, the combination of SERS and machine learning has shown immense promise as a viable characterization technique due to its extreme sensitivity[32, 33], with recent applications ranging from cancer[31, 34] to infectious[35, 36] and reproductive diseases[37].

Raman spectroscopy relies on the nonlinear interaction of incident light and the investigated sample, where the sample essentially acts as a dispersive medium, and the resulting inelastically scattered light is collected by a set of optical components. The light is then decomposed using a grating and a charge-coupled device (CCD) to measure the specific intensity of each frequency component[29]. The spectral range of modern CCDs can range from several hundred to several thousand pixels and is based on the properties of the source light, grating, and other involved optical components. Usually, the acquired spectra contain thousands of pixels, providing a unique, high-resolution “fingerprint” of the investigated sample based on the relative shape, position, and intensity of the peaks.

While characterizing specific spectral changes in chemistry applications may be possible by tracking basic changes in single or multiple peaks, the investigation of more complex samples as in most biomedical applications requires multivariate analysis tools such as Principal Component Analysis (PCA) [38] followed by other machine learning applications [31, 35, 39, 40]. In Euclidean space, a coordinate system is defined by the resolution of the CCD, and each pixel gives an independent value that can be considered as one dimension of the spectral data. PCA finds a linear transformation (a rotation of the coordinate system) that maximizes the variation of the data in the new principal axis, with the first principal axis representing the maximum variance, followed by the second, and so on. The first two axes of PCA and the projection of spectra within them are commonly used to show the separation of the data or the formation of clusters. The PCA axes therefore, indicate all the CCD’s pixels (i.e. all the possible peaks in the dataset) with different weights or PCA scores assigned to each pixel. This ultimately enables the quantitative interpretation of the separation of the data in the PCA space and the correlation of automatic clustering to a specific peak or peaks.

Another application of PCA involves denoising the acquired Raman spectra[31, 35, 41]. As the variance of the data is maximized within a few principal axes, the higher principal axes carry little to no information about the data set. Therefore, after applying the PCA, one can consider the first few axes as a new set of features for the data instead of the CCD’s values. Then, using the calculated PCA scores of each principal axis, the reverse transformation (from PCA to CCD’s axes) is possible to obtain the denoised spectra. The same technique has also been used for dimensionality reduction, where the first PCA axes can be considered as a new set of features that represent most of the spectral information, and can be subsequently used by automated machine learning models.

However, PCA has several major limitations associated with the linear nature of its transformation. For example, if the spectra are located on a nonlinear manifold in the Euclidean space associated with the CCD’s pixels, the PCA cannot effectively perform dimensionality reduction or data visualization. While there are other dimensionality reduction methods such as t-SNE [42] and UMAP [43, 44] that find and unfold nonlinear manifolds in their projection space using nonlinear transformations, their projection space lacks interpretability, and the reasons behind the formation of the clusters remains unknown. Another PCA shortcoming is the high number of principal components required to reconstruct the data from its projection space, often requiring more than a dozen principal components to reconstruct more than ninety percent of the information. This is again due to the linear reconstruction of the spectra from their principal projections such that the information in the selected principal axes are limited to a linear combination of their associated PCA scores. These limitations indicate the need for more advanced tools, such as deep learning, to take full advantage of the highly complex and information-rich Raman spectra from biological samples.

Autoencoders are a subtype of deep learning designed to extract the latent features from unlabeled datasets [25, 45–50]. They accomplish this by first using an encoder to compress the dimensionality of the data into a lower dimensional space known as latent or code space, and then restoring it back to original dimensions by reconstructing the input at the output. Many autoencoder variations have been introduced for different purposes such as dimensionality reduction [45], denoising of images, preprocessing of spectroscopy data [25, 46–48], generative models [49], and data storage compression [50]. However, despite their potential in allowing quantitative analysis of highly heterogeneous and dynamic Raman spectral data sets, few studies have applied autoencoders to the study of EV SERS to date.

In this work, we combine SERS with a novel architecture of a hybrid autoencoder to enable the label-free quantification of the percentage of FBS-derived EVs within an EV mixture. The proposed network linearly compresses EV Raman spectra and uses a nonlinear decoder to reconstruct the data from the linearly embedded information. However, unlike conventional autoencoders, we also connect the code (latent) layer to an additional output that forces the encoder to reconstruct the data while also considering the data’s unique labels (known EV ratios). This network is easily tunable and can range from unsupervised to supervised by simply changing the weight of the reconstructing decoder relative to the data’s labels. This approach was first tested using known mixtures of chemical standards, then used to quantitatively detect the relative amount of FBS-derived EVs within a mixture of breast cancer EVs from a bioreactor culture. Surprisingly, the network could also be used for knowledge discovery, where fitting a polynomial curve to the code (latent) space values of Raman spectra from a chemical milling process allowed accurate prediction of the spectra from unknown portions of the reaction. Based on these results, this powerful method should advance the study of EVs in diagnostics, biomanufacturing, and therapeutics, while also enabling improved analysis in many other Raman spectroscopy applications.

## 1 Methodology

### 1.1. Neural Network

Consider the network architecture depicted in Supplementary Figure 1. At the beginning of the network, there is a stack of dense layers followed by linear activation layers. This first section is known as the encoder because each consecutive layer gradually compresses the high dimensional Raman spectra at the input into a lower dimension layer known as the latent or code layer.

To construct the combined decoder, we consider two parallel networks. In decoder A, there is a stack of dense layers with an ascending number of dimensions to eventually match its output layer dimensions to that of the input data. For decoder B, there are several stacks of dense layers (in parallel to decoder A) that match its output to the data’s corresponding output, which can be either the data’s labels in the case of classification, or any scaler number associated to each data set in the case of a regression.

#### 1.1.1 Encoder

Liner activation was used in the encoder’s layers to ensure that the output of each node is simply a linear combination of its input plus a bias, as in equation **2 and Supplementary Figure 1. In this equation**, 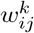 are the weights associated with each node at the previous layer where indices *k, i*, and, *j* denote the layer, the node that is being calculated, and the previous layer’s nodes, respectively 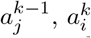 and, 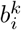 are the outputs of the previous layer’s nodes, outputs of the current layer’s nodes, and the bios associated to each node at the current layer, respectively.

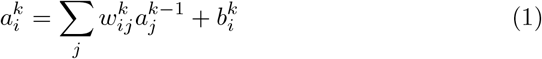

Therefore, without any compensation on the generality of the encoder layers, the output of the latent space can be written as equation 2, in which the *a*^*latent*^ and *x*_*j*_ are the components of the latent and input data while 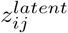 and 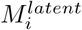 are the equivalent weights and biases that can be found based on equation 1.

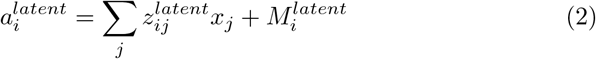

Please note that the weight associated with each node at the latent layer 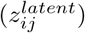 can be considered as a vector with the same dimension as the input vector (*x*_*j*_). In other words, the latent space (defined by the output of the latent layer) is a linear, lower-dimensional projection of the data with some biases. This lower dimensional subspace is defined by the vectors 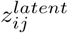 and biases 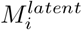. For example, in the case of the latent dimension equal to two, the latent space is a plane indicated by two vectors 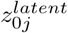 and 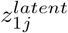 As a result, the projected points of the input data are interpretable and their positions in the latent space directly indicate the absence or presence of 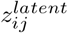vectors in the original data.

Therefore, the selection of the latent space is done by selecting the proper vectors 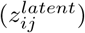 and biases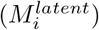, based on the defined goal and the loss function at the end of the decoders’ layers. In deep neural networks, it is also common to use batch normalization immediately following each dense or convolution layer. These added layers are also linear transformations, and by applying them we maintain all the advantages of the mentioned linear activation while also regularizing the model and reducing its sensitivity to the initialization [51] (for the case of the decoder).

#### 1.1.2 Decoder

Decoder A - In order to extract the highly nonlinear and complex information from the linear projection of data of the encoder, multiple dense layers are used with rectified linear unit (ReLU) activation. Also, to avoid problems such as accuracy saturation and gradient vanishing, some skips similar to the those in ResNet are also used. The decoder results in data with the same dimension as the input and attempts to recreate the input with minimal error. Importantly, the linear encoder always produces a simple and explainable latent space, while the decoder always retrieves as much information as possible from the latent space, regardless of the complexity required to do so.

Decoder B - Several non-linear activated dense layers are used to match the data’s labels or any corresponding numerical output, allowing this portion of the decoder to act as a supervised classifier or regression. A similar network has been previously introduced as a bottleneck classifier for simultaneous dimension reduction and classification [37].

#### 1.1.3 Network Implementation and Training

The proposed network was implemented via the stabilized TensorFlow package [52] in Python. Adam optimizer as a variant of stochastic gradient descending is used to train the network while learning rates, *β*_1_, *β*_2_ and *ϵ* were chosen to be 10^*−*3^, 0.9, 0.999, and 10^*−*8^, respectively.

### 1.2 Chemical Standard Mixtures

Solutions of Rhodamine 6G and Rhodamine B were prepared in ultrapure water at a concentration of 10^*−*6^M. These solutions were carefully mixed to produce different volumetric/molar ratios, including 100-0, 99-1, 95-5, 90-10, 83-17, 80-20, 75-25, 65-35, 50-50, 35-65, 25-75, 20-80, 17-83, 10-90, 5-95, 1-99, 0-100, followed by drying them on a thermally evaporated gold thin film on a standard glass slide. Raman measurements were performed using Horiba LabRam Evolution confocal Raman microscope with a 532nm laser, 50× microscope object, and 1 percent of the maximum laser power which is 100 mw. The acquisition time for each spectrum was also chosen to be 1 second, and 100 spectra were collected over a 10 × 10 micron rectangular grid for each chemical mixture, resulting in 1700 spectra for this dataset.

### 1.3 Extracellular Vesicle Mixtures

Spike-in mixtures of two distinct small EV populations were produced by first isolating EVs from commercial FBS (Sigma, F8067 Batch 16A327) and conditioned media from MDA-MB-231 cells cultured in a CELLine AD1000 bioreactor with chemically defined serum replacement CDM-HD (Fibercell Systems) as previously reported and thoroughly validated according to MISEV 2018 guidelines[53–55]. Importantly, we refer to FBS-derived EVs and particles (EVPs) as simply “FBS EVs” throughout the text to focus on the analysis approach, but fully acknowledge that several other types of particles, including lipoproteins and growth factor aggregates are co-isolated using our reported workflow. These non-EV contaminant particles actually demonstrate the robust nature of label-free SERS combined with this analytical approach, as highly heterogeneous spectra can still be used for robust quantification. Additionally, a small amount of FBS was used in the upper bioreactor chamber throughout the MDA-MB-231 culture such that small molecules (¡10 kDa) could pass through the semi-permeable dialysis membrane and interact with the MDA-MB-231 cells and EVs while the FBS EVs were completely excluded from the cell chamber and the conditioned media[56]. Thus, these two EV samples are ideal, albeit simplified models that reflect unknown EV mixtures from conventional cell cultures. FBS was first diluted 1:1 in Advanced DMEM/F12 (ThermoFisher 12634010), the same media used in the bioreactor cultures. In all cases, EVs were isolated using differential ultracentrifugation (10,000 xg for 30 min, then 100,000 xg for 70 min, Beckman Avanti J-30I) followed by size exclusion chromatography (qEV Original, Izon) and ultrafiltration (100 kDa Vivaspin 500, Sartorious) to exchange from PBS buffer to ultrapure water. EV concentrations were then measured using nanoparticle tracking analysis (NTA, Nanosight NS300) with the same settings for each sample (recording - 3 × 30 second videos, screen gain 1, camera level 14; analysis - screen gain 10, detection threshold 6) and normalized to 2.6*x*10^11^ particles/ml by diluting in ultapure water immediately before SERS experiments. An EV production and isolation workflow, NTA data of the isolated EVs, and negative staining TEM images of the MDA-MB-231 bioreactor-derived small EVs (performed as described in [54]) are shown in Supplementary Figure 2.

For comparison, the same FBS (Sigma, F8067 Batch 16A327) was stored in three basal media formulations (ThermoFisher DMEM 11995065, Adv DMEM 12491015, and Adv DMEM/F12 12634010) for 2 weeks at 4 C prior to performing the EV isolation and characterization workflow. This was done to determine whether storage in the basal media formulation could affect the final FBS EV Raman spectra. Finally, FBS EVs from three manufacturers (Sigma F8067-500ML Batch 16A327, Moregate MG-FBS0820-500ML-GEN Batch 55827102, Gibco 10091148 Batch 2282152P) were also isolated and normalized using an identical workflow and compared using SERS to determine whether FBS EVs from different suppliers produced classifiably distinct Raman spectra.

EVs are known to be highly heterogeneous in their biochemical cargo, even from single cell lines, and thus their Raman spectra are also very complex. Additionally, their small volumetric sizes do not allow straightforward acquisition of Raman spectra using spontaneous Raman techniques. Therefore, we again used the femtosecond laser-fabricated LINST SERS substrate to enhance the EV Raman signal[37]. EV mixture ratios of 100-0, 95-5, 90-10, 80-20, 50-50,

80-20, 90-10, 95-5, and finally 100-0 of FBS and MDA-MB-231 EVs were prepared, and 20 microliters of EVs from each mixture compositions were pipetted onto the LINST surface and dried at 40 C in a drying oven. Then, 1156 spectra were obtained over a grid of 34 × 34 using a 50 × microscope objective and a 100 mW, 785 nm laser attenuated to 25% of its maximum power. Acquisition time of 100ms (.1s) for each spectrum was used so that the total acquisition time for all 1156 spectra did not exceed 30 minutes. This is important as a long signal acquisition time can lead to sample degradation and thereby unwanted change in the spectra of the EVs. We then denoised the spectra and removed their baselines using Asymmetric Least Square smoothing (AsLS) and Savitzky Golay filters, respectively. We also divided 1156 spectra of each sample into 76 batches containing around 15 spectra each and calculated the average of the spectra for each batch for use in subsequent analyses. This averaging technique reduces the heterogeneity of the spectra in each batch as the averaged spectra should contain more diverse peaks than any single EVs spectra while reducing the total number of spectra used for each sample from 1156 to 72.

## 2 Results and Discussion

To demonstrate the utility of the presented network for various problems, we applied it to multiple synthesized data sets, real-time and in-situ Raman spectroscopy of a chemical reaction, Raman spectra from a chemical mixture, as well as Raman spectra from an EV mixture of bioreactor-produced MDA-MB-231 and Sigma FBS small EVs.

### 2.1 Synthesized Data

#### 2.1.1 Univariate Data

In the first example, we considered continuous shifts of an identical Gaussian peaks from the left of the spectral range to the right as shown in Figure 1 (a) and Equation 3.

**Fig. 1.**
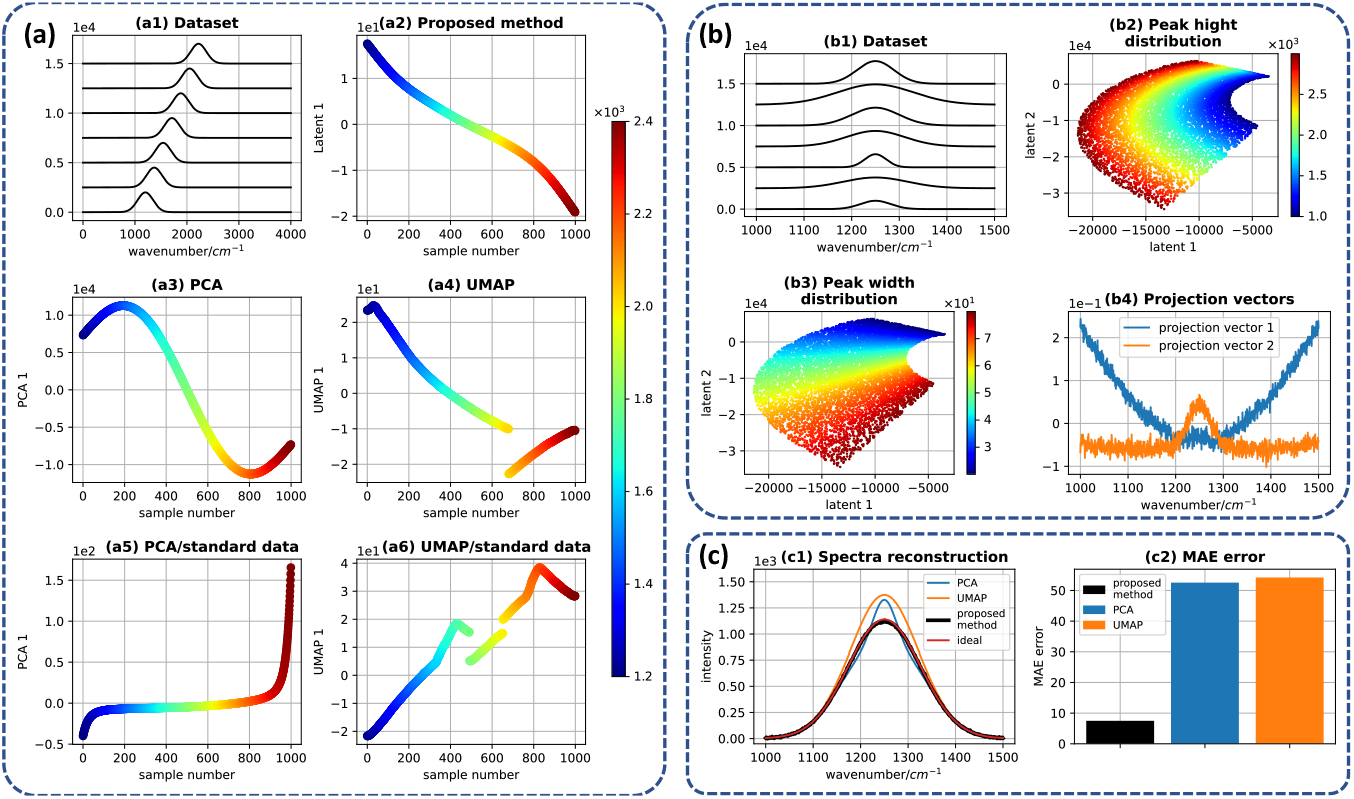
(a) Synthetic dataset example with one independent parameter. (a1) Synthetic spectral data with a continuous shift of the peak position of an identical Gaussian peak, (a2) one-dimensional unsupervised projection of the dataset using our proposed method. (a3) and (a4) one-dimensional PCA and UMAP projections of the dataset when the dataset is not standardized, respectively. (a5) and (a6) one-dimensional projection of the dataset when the dataset is standardized, respectively. All the color bar in (a) shows the peak position of each spectrum in the dataset. (b) Synthetic dataset example with two independent parameters, (b2) and (b3) show the two-dimensional projection of the dataset while the color of each point shows the peak and width distribution of each projected spectrum, respectively. (b4) The linear projection vectors obtained by the proposed method.

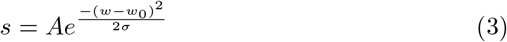

In this equation, *s, A, w, w*_0_ and *σ* denote the spectra, amplitude, spectral number, center of the peak, and its variance, respectively. We know that each synthesized spectrum, in this example, can be uniquely identified by its peak positions. Moreover, as the equation 3 is continuous and infinitely derivable with respect to *w*_0_, the continuous change of the peak centers creates a continuous Riemannian one-dimensional manifold in the high dimensional space denoted by the given spectra. Therefore, an ideal dimensionality reduction method should be able to compress all the information of any of the spectra to a unique, single number. As a result, the latent projection of the continuously varying data depicted in Figure 1 (a) should create a strictly increasing or decreasing trend.

The one-dimensional projection of the described data using our proposed method when the decoder is chosen to be just A is depicted in Figure 1 (b) Similar results for both one-dimensional PCA and UMAP projections when the given data is either standard scaled or not are also shown in Figure (c)-(f). In these figures, each spectrum is depicted in a 2D plane while the horizontal axis is the sample number associated with each spectrum in the data set, while the vertical axis shows the projection result for each spectrum, and the color of the line combined with the color bar shows the peak position of each spectrum. Additionally, the standard scaler transformation linearly projects the value of each spectrum at each wavenumber so that the entire data set has a mean of 0 and standard deviation of 1 at any given wavenumber. This standard scaler transformation is a routine practice before using PCA and UMAP to ensure that each wavenumber plays an equal role in its given projections.

Clearly, neither PCA nor UMAP produced strictly increasing or decreasing trends in their embedding space when dealing with non-standard data. As such, both methods projected completely different spectra within the data set onto identical points within the embedding space. Therefore, they likely need more than one dimension in their embedding space to accurately represent the data. However, the PCA was more successful in achieving a continuous trend, while UMAP produced one discontinuity in its latent projection. Furthermore, while PCA was more successful when fed with standard scaled data, UMAP again resulted in a non-continuous trend.

The resulting discontinuity in the UMAP projection is likely due to a nonlinear selection of the one-dimensional manifold and its dependence on the choice of “number of neighbors” within its algorithm. This number should be manually selected and can significantly affect the projection result. Such discontinuities in UMAP results are also very problematic for the interpretation of the data using its embedding, because two spectra that are actually very similar may appear at the two ends of the discontinuity. In clear contrast, the proposed method results in a continuous, strictly increasing trend. Therefore, each spectrum is projected onto a unique point in the embedding space, which is the ideal case for representing this data in 1D space.

#### 2.1.2 Multivariate Data

For the second example, spectra were synthesized with the same peak positions but randomly selected peak widths (*σ*) and heights (*A*) in Equation 3, using peak heights between 1000 to 3000 and peak widths between 20 to 80. Again, continuous changes of these two variables in Equation 3 should result in a two-dimensional Riemannian manifold. This data is then projected into a two-dimensional latent space resulting from our proposed method similar to the previous univariate problem. Some spectra of this data set and its projection in a two-dimensional latent space with both width and height distribution are depicted in Figure 1 (b1), (b2), and (b3), respectively. Clearly, each spectrum can be distinguished with different peak heights and widths in a two-dimensional representation of the data. Interestingly, spectra with similar heights and widths are distributed along two different axes of the embedding shape. Finally, the projection vectors 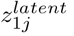 and 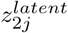 calculated from the eights and biases of the linear encoder are shown in figure 1 (b4).

Another major advantage of the proposed method is its outstanding ability to reconstruct the projected spectra based on their embedding information compared to other techniques such as PCA and UMAP. To show this, we projected all of the spectra of this example into a two-dimensional space using PCA and UMAP and did the reverse transform to assess the reconstructed spectra. The output of the decoder A for each spectrum was also used for this comparison. In Figure 1 (c1), the reconstructions of one of the spectra of this data set using PCA, UMAP, and our method are presented. The mean absolute error of all reconstructed spectra based on the ideal form value (the input spectra in this example) were also calculated and are depicted as a bar chart in Figure 1 (c2). These results clearly demonstrate the superiority of the presented technique compared to the other methods in preserving the information using its latent projection. The main reason for this ability is the complex network architecture used for decoder A. Such deep and complex architectures should be capable of extracting the complex distribution of the spectra and their patterns in the latent space and converting them to the actual input data at its output.

#### 2.1.3 Generalization and Gap Filling

To compare the generalization of the problem using these three distinct techniques, a patch of the data was removed (spectra with a height between 1700 to 2300 and width between 40 to 60) and used as the test set, and the remaining data was used as the training set. Then, all methods were trained using only the training set to project the data in 2D space. These training data sets were expected to result in a 2D manifold but with a hole in its middle, which was clearly the case as shown in Figure 2 (a1), (b1) and (c1) for our proposed method, PCA, and UMAP, respectively. Then, the testing set was projected using the trained network. In an ideal case, the projection of the test set using each method should effectively fill its corresponding hole in Figure 2 (a1), (b1), and (c1). The projected test set using the proposed method, PCA, and UMAP are shown in Figure 2 (a2), (b2), and (c2), respectively, and the overlaid projection of both the training and test sets are shown in Figure 2 (a3), (b3), and (c3). Interestingly, only the proposed method and PCA projected the testing set in such a way that training set projection gap is filled. This is thanks to the linear nature of dimensionality reduction used in the proposed method and PCA, which allows them to preserve the continuity of the high-dimensional data within all lower-dimensional projections.

**Fig. 2.**
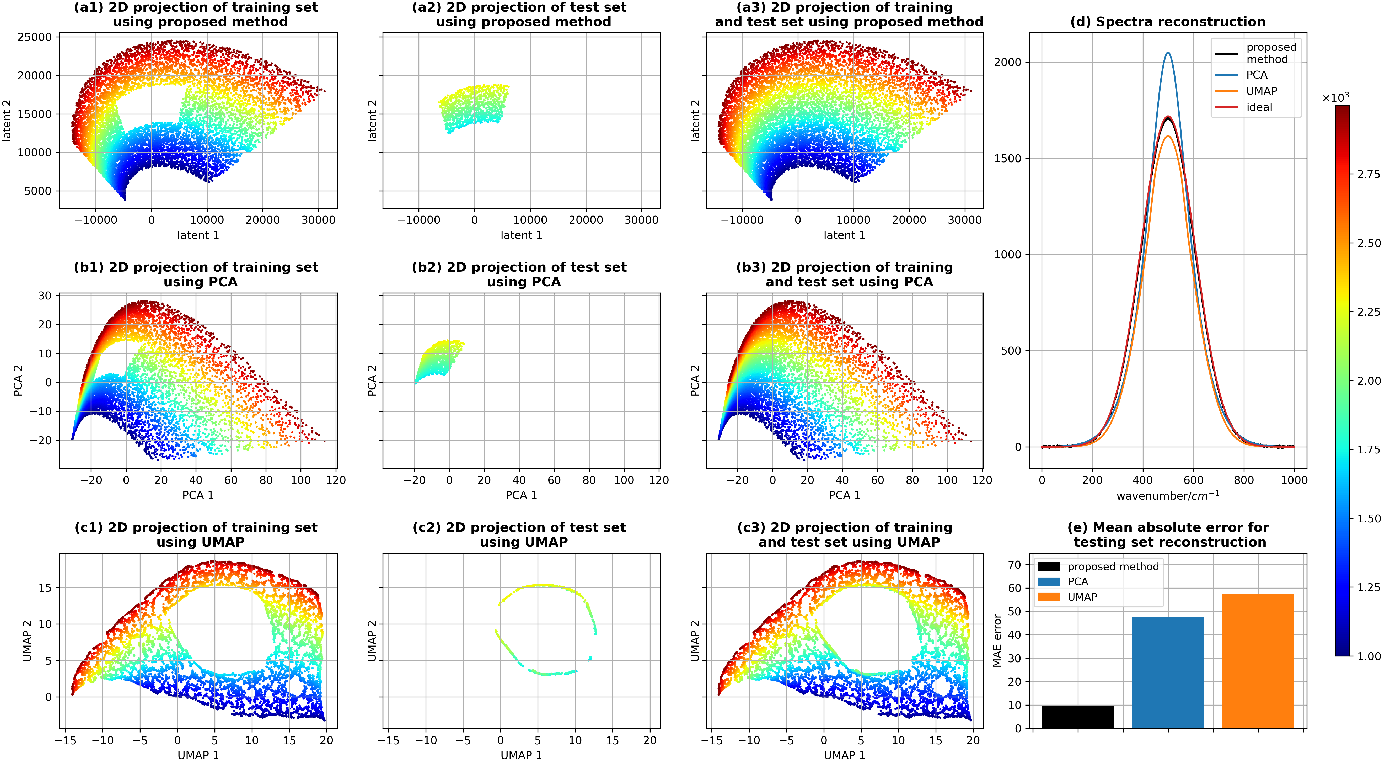
(a1), (b1), and (c1) The two-dimensional projection of the training set using our unsupervised method, PCA and UMAP, respectively. (a2), (b2), and (c2) The two-dimensional projection of the testing set using our proposed trained network, PCA, and UMAP, respectively. (a3), (b3), and (c3) Overlaid projection of both training and testing sets for our method, PCA, and UMAP, respectively, where the color of each point shows the peak height of the corresponding spectrum in the dataset. (d) One of the reconstructed spectra of the testing set using our proposed method,PCA, and UMAP vs the ideal output. (e) The MAE error of the reconstructed spectra for the testing set for our proposed method, PCA, and UMAP.

Additionally, we attempted to reconstruct the testing set spectra from its projected information using inverse transform to see which method is ideal for reconstruction. Similar to the previous example, we used the mean absolute error between the reconstructed and the ideal spectra to determine their relative performance. One of the reconstructed spectra using each of the various methods, as well as its ideal shape, is presented in Figure 2 (d), while the mean absolute error is depicted in Figure 2 (e). Please note that using our proposed method, spectra are reconstructed in decoder A only. Clearly, while this part of the network is complex, it generalized the problem well and did not become overfitted. Thus, such a decoder may be used to predict the unknown data within the gap or potentially even fill the gap during the data acquisition stage. A clear example of this approach is demonstrated in the next section.

### 2.2 Chemical Reaction

In [17], researchers showed that Raman spectroscopy is a potential candidate for in situ monitoring of chemical milling reactions. They demonstrated this by obtaining a series of Raman spectra from a 1:1 stoichiometric mixture of nicotinamide (Nic) (a form of vitamin *B*_3_ used in vitamin *B*_3_ complex medication) and salicylic acid (Sal) (as an active form of acetylsalicylic acid and an ingredient of antacid drug Pepto-Bismol[17]) over 1 hour with the time resolution of 15 seconds. Over time, the Raman spectra of the sample changes simply due to the formation of the Nic and Sal co-crystal.

Regardless of the number of intermediate products or the complexity of the reaction, “time” (from the beginning of the reaction) can identify a unique state of the reaction as long as the other parameters (such as temperature or pressure) are controlled. Therefore, we expect the Raman spectra available in their dataset to form a smooth one-dimensional manifold. All Raman spectra obtained in this study are shown in Figure 3 for Raman shift between (0-2184*cm*^*−*1^) and (300-1800*cm*^*−*1^). The spectral band between (300-1800*cm*^*−*1^) is depicted for the sake of better visualization of the changes of the Raman peaks in this band and all multivariate analyses were performed on the whole spectral band.

**Fig. 3.**
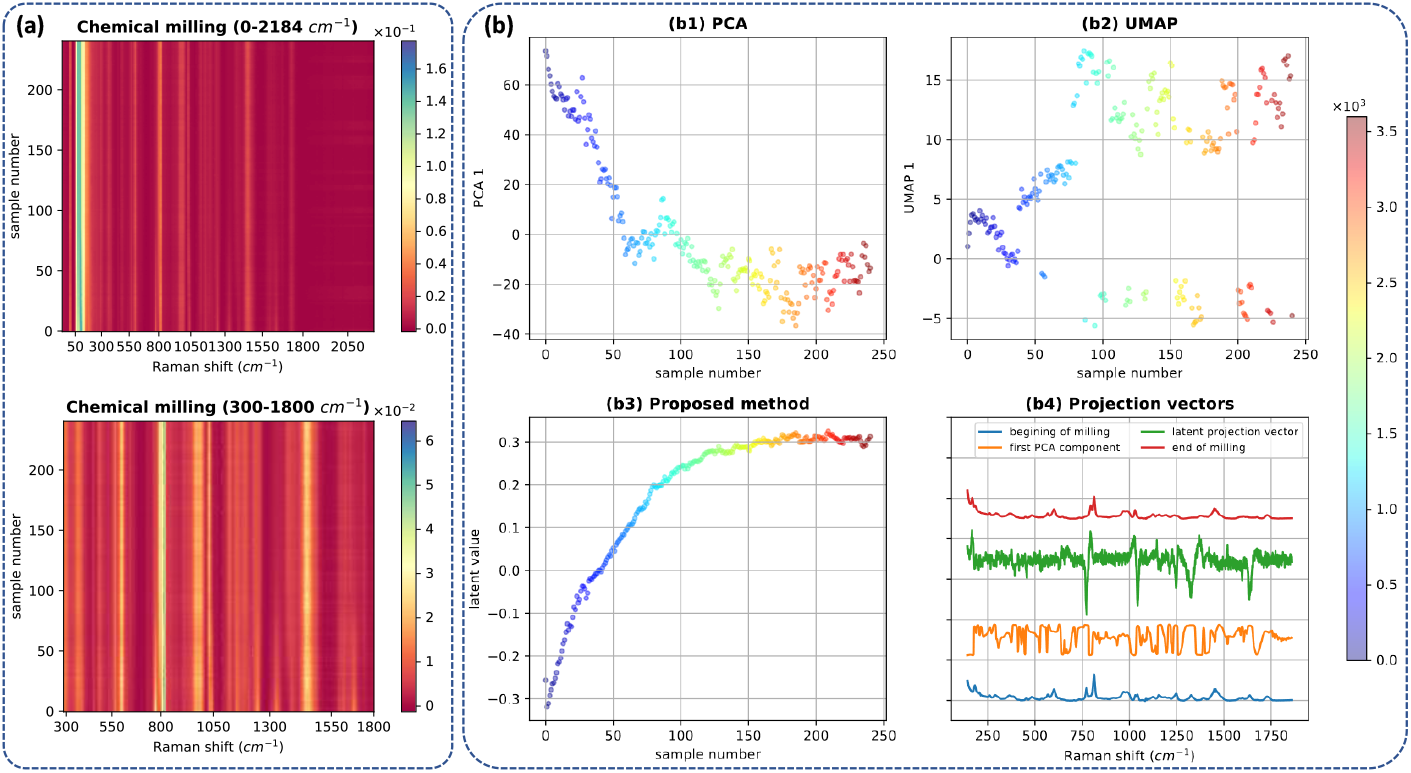
(a) Raman spectra of chemical milling reaction for spectral width of (0-2184*cm*^*−*1^) and (300-1800*cm*^*−*1^). Unsupervised one-dimensional projection of all spectra in latent space of PCA (b1), UMAP (b2), and the proposed method (b3). Raman spectra at the beginning and end of the reaction, as well as the calculated projection vector using our proposed method and PCA (b4).

PCA, UMAP, and the projection result from our method are depicted in Figure 3. Please note that the data were standardized according to the method described in Figure 1 before applying both PCA and UMAP. In these graphs, the vertical axes show the projection values from each method, while the horizontal axes and color of each point show the sample number and time from the beginning of the reaction in seconds, respectively. Again, the proposed method clearly outperforms both PCA and UMAP in describing the nature of the process using a 1D projection, as both PCA and UMAP could not achieve a strictly increasing or decreasing trend. The other problem of PCA and UMAP is that for any given time point in the reaction, the spectra are projected to a band in the projection space and cannot be assigned to specific times. In the proposed method, projection values can be correlated to their corresponding time points in the reaction. This is especially true from the beginning of the reaction and until it reaches steady state.

#### 2.2.1 Gap Filling in a Chemical Reaction

To test the ability of the proposed network to fill the gap within a dataset, we purposely removed the spectra between (20-60 sample numbers, i.e. between 300-900s from the start of the reaction) and used the rest as the training set as shown in Figure 4 (a1)-(a3). Then the training set (Figure 4 (a2)) was used to train the network and resulted in the projected points depicted in Figure 4 (b1). Due to the application of the linear encoder, the gap also appeared in the projection of the points. Now, the gap can be filled by first fitting a polynomial curve onto the projection and sampling from the gap (as shown in Figure 4 (b2)). Finally, if the sampled points are passed through the decoder only, one can reconstruct the spectra from the gap region. Generated spectra from the decoder and the actual spectra of this gap are depicted in Figure (d1)-(d3) and (e1)-(e3), respectively. Clearly, all of the peaks appear in the correct locations, and the trend and rate of the changing of the peaks through time clearly match. In effect, we can conclude that the decoder has essentially learned the unique mechanism of this specific reaction based solely on its inelastic scattering.

**Fig. 4.**
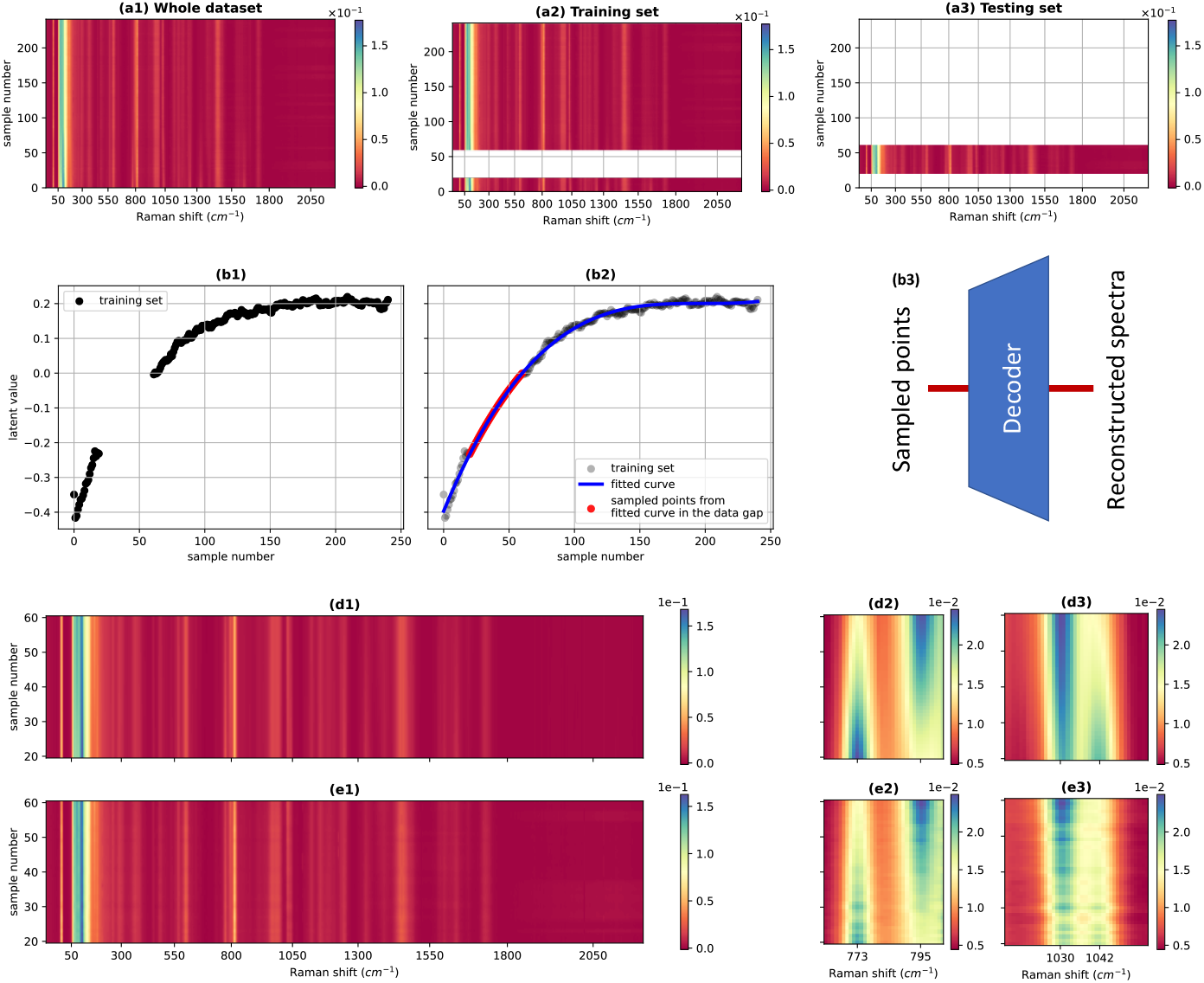
(a1), (a2), and (a3) whole, training, and testing datasets. (b1) one-dimensional latent projection of the training set using our method. (b2) curve fitting followed by sampling the gap area. (b3) passing the sampled points into the decoder to construct missing data. (d1), (d2), (d3) the constructed missing spectra by the decoder. (e1), (e2), and (e3) the testing set (missing data).

9 m

### 2.3 Raman Spectra of a Chemical Mixture

All the baseline corrected, denoised, and Euclidian-normalized spectra for R6G and R6B mixtures are shown in Figure 5 (a). The unsupervised one-dimensional projection of all acquired spectra using PCA, UMAP, and the proposed method, when decoder A is selected, are shown in Figure 5 (b1), (b2), and (b3), respectively. The reason that we again chose the one-dimensional projection is that the change of the Raman spectra when we smoothly change the mixture ratio should also be smooth and only depend on a single number, the ratio of two chemicals. Again, it can be seen that the proposed autoencoder achieved better results in describing the trend of the data. The projection vector, 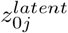 of the latent space and the average Raman spectra pure Rhodamine 6G and 6B are also shown in Figure 5 (b4). Interestingly, the point of any changes between R6G and R6B spectra are identified by the latent projection vector of the proposed method.

**Fig. 5.**
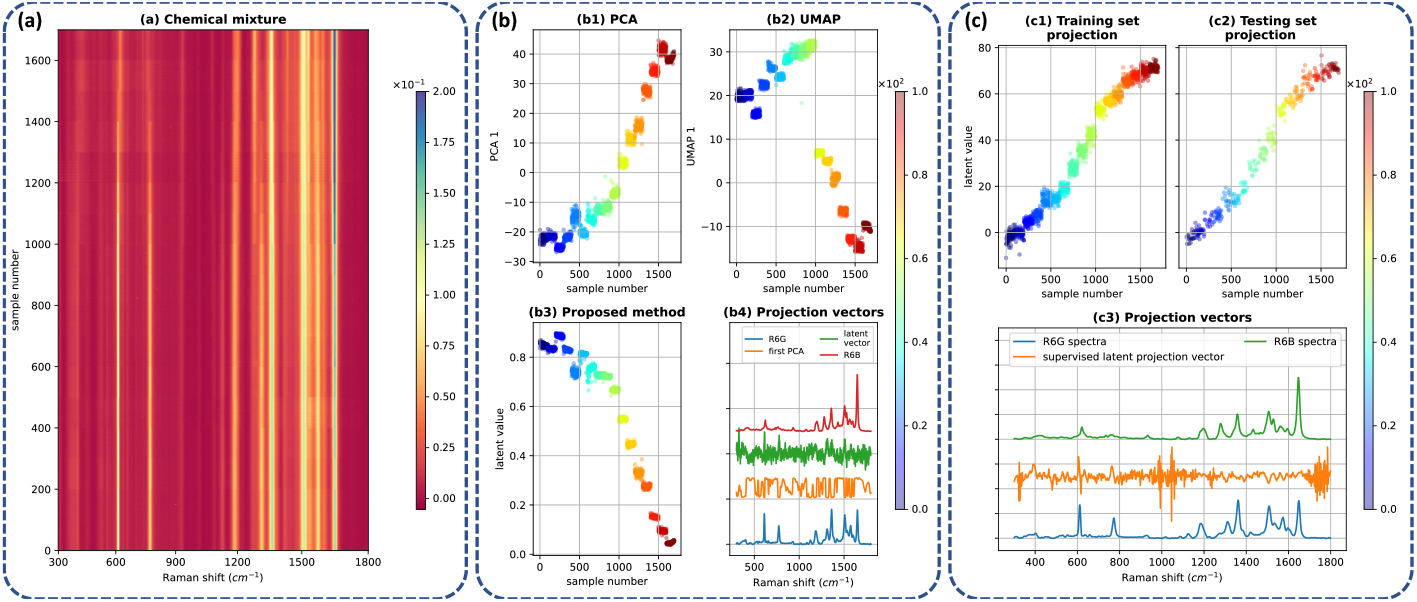
(a) Raman spectra obtained from various concentrations of R6G and R6B. (b1), (b2), and (b3) Unsupervised one-dimensional projection of all spectra using PCA, UMAP, and our proposed method, respectively. (b4) The projection vector obtained by the PCA and our proposed method, and the SERS Raman spectra of pure R6G and R6B. (c1) and (c2) Supervised one-dimensional latent projection of the training and testing sets, respectively, using our proposed method. (c3) The calculated supervised projection vector and Raman spectra of pure R6G and R6B.

For this example, we can also incorporate decoder B with decoder A simultaneously. We selected the decoder B to act as a regression and determine the percentage of R6G in the mixture. In such cases, the errors generated from both decoders are backpropagated through the encoder, which finds a more descriptive latent projection vector that both reconstructs the spectra and determines the percentage of the mixture.

### 2.4 Extracellular Vesicle Mixture

The resulting preprocessed Raman spectra from the EV mixtures, both before and after averaging, are presented in Supplementary Figure 3. KMeans clusters with their corresponding bar graphs showing relative cluster abundance for SERS data from EV mixtures with a cluster number of k=8 are plotted in Figure 6. Based on the distribution of labels 6 (a), the heterogeneity of labels for each sample’s spectra gradually reduce from 100-0 (pure FBS EVs) to 0-100 (pure MDA-MB-231 EVs). This was expected as the FBS EVs are derived from fetal bovine serum and naturally contain thousands of EV subtypes from different tissues, while the breast cancer EVs are derived from a single cell line. Importantly, the FBS EV sample undoubtedly contains large, unknown quantities of lipoproteins and other nanoparticle contaminants, demonstrating the robust nature of the network design. KMeans clustering with cluster number of (k=2) is also represented in Figure 6 (b). This graph clearly shows that progressing from pure FBS EV to pure MDA-MB-231 EVs results in the population of one cluster (purple) decreasing and the other cluster (turquoise) increasing. This indicates that the cluster center associated with the purple is more related to the spectra of the FBS EVs while the turquoise cluster is more related to the spectra of the MDA-MB-231 EVs.

**Fig. 6.**
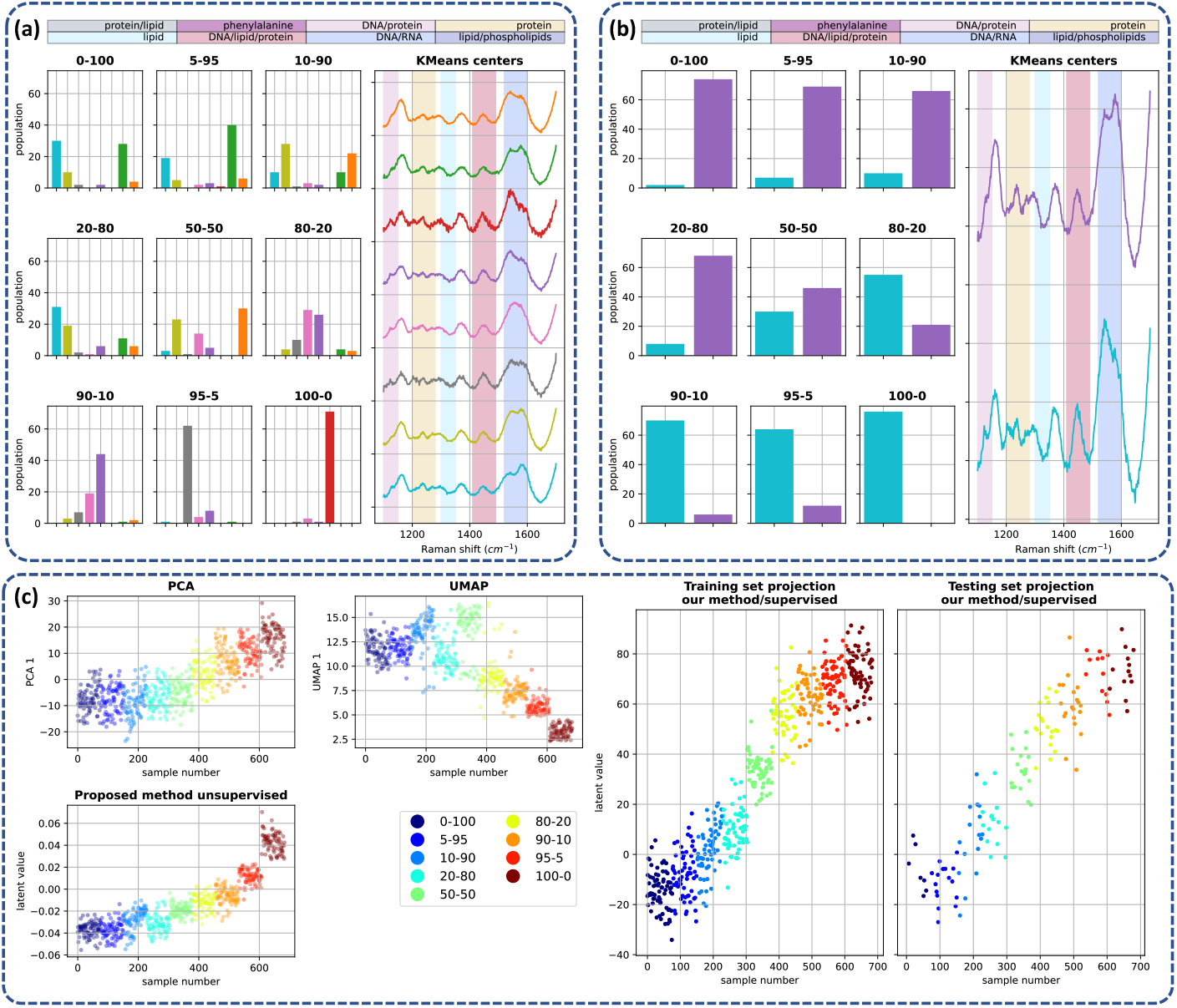
(a) Label distributions in KMeans clustering when K is equal to 8 for each composition of EV mixture. Clusters’ centers are also depicted with the same color as their associated bar in each composition. (b) Label distribution and their associated center when K is equal to 2. (c) Unsupervised projection of the EV SERS spectra using PCA, UMAP, and our method. The color of each projected spectrum indicates the percent of the EV composition. Supervised projection of the training and test sets using our proposed method.

This dataset was fed into our proposed autoencoder considering only decoder A for the purpose of unsupervised dimensionality reduction. We also performed one-dimensional PCA and UMAP transformations of this data as shown in Figure 6 (c). Similar to the previous example, we can see that the trend obtained by the proposed method is more compact in our proposed method while showing an increasing trend in its latent projection. EVs derived from FBS from different suppliers were compared using the same EV workflow, with no apparent differences in spectra produced from Gibco, Moregate, and Sigma FBS-derived EVs as shown in Supplementary Figure 4. Sigma FBS-derived EVs that were stored in either Advanced DMEM/F12, Advanced DMEM, or DMEM at 4C for 2 weeks were also compared using an identical workflow, with no apparent differences as shown in Supplementary Figure 5.

Another interesting result achieved by both UMAP and the proposed dimensionality reduction techniques (shown in Figure 6 (c)) is that, the spectra for MDA-MB-231 EVs were clustered completely separately from the other samples. There are many potential reasons that explain this phenomenon. First, FBS EVs are smaller in size, which likely allows them to fit more easily within the SERS hotspots. This may cause them to provide a more substantial contribution to the SERS spectra of the composition. Another reason is that in our averaging process, the samples that contain FBS EVs were averaged and therefore no pure spectra of breast cancer remained in their spectral data set. As there are no FBS EVs in 0-100 sample, it likely lacks the peaks associated with the FBS EVs and is more effectively separated.

To have a better separation between the different compositions we also incorporated the decoder B as a regression unit and performed supervised training with known mixture compositions. This decoder is utilized with a much smaller loss weight (10^−3^) compared to decoder A (when the loss weight of decoder A is considered to be 1). This forces the network to first learn to set its weights and biases in a way that accomplishes the unsupervised task (spectral reconstruction). Once the loss of decoder A is small enough, decoder B then fine-tunes the network parameters to achieve a good regression result and obtain the composition mixture. Please note that the resulting representation of the spectra in the latent layer should still be sufficient to reconstruct the entire data set at the output of the A decoder. Otherwise, it will be heavily penalized due to its much larger loss weight. Also, as we are incorporating supervised learning, the EV mixture data set was divided into training and test sets.

## 3 Conclusions

In this work, we propose a novel deep learning framework based on a custom autoencoder architecture for interpretable dimensionality reduction and data analysis in Raman spectroscopy applications. This network successfully completed complex tasks including the interpretable analysis of dynamic chemical reactions, as well as EV and chemical mixtures based solely on the acquired Raman spectra. We also demonstrate that this manifold learning approach is superior to conventional techniques such as PCA and UMAP while offering a high level of interpretability for both supervised and unsupervised learning. Due to the linear encoder used in this method, the network can even fill a gap within a Raman spectral dataset of a chemical milling reaction, suggesting its applicability to many other Raman spectroscopy applications.

## Author Contributions

MK: conceptualization, data curation, formal analysis, investigation, methodology, software, validation, investigation, visualization, writing. MM: conceptualization, methodology, writing. RO: investigation. AA: investigation. MML: investigation. XN: writing. ER: writing. ZDS: writing. WX: resources, supervision, project administration, funding acquisition. CB: writing, funding acquisition, resources. LWC: writing, funding acquisition, resources. NGRB: resources, supervision, project administration, funding acquisition. CLH: conceptualization, data curation, methodology, validation, investigation, resources, writing, supervision, project administration.

## Acknowledgements

The authors would like to thank the Breast Cancer Foundation New Zealand, University of Auckland’s Hub for Extracellular Vesicle Investigations, Ohio State University’s LEGACY Postdoctoral Scholars program, and the NIH (R01 GM109988, K99EB033857) for their continued support.

