## Supplementary Figures for "Manifold Learning Enables Interpretable Analysis of Raman Spectra from Extracellular Vesicle and Other Mixtures"

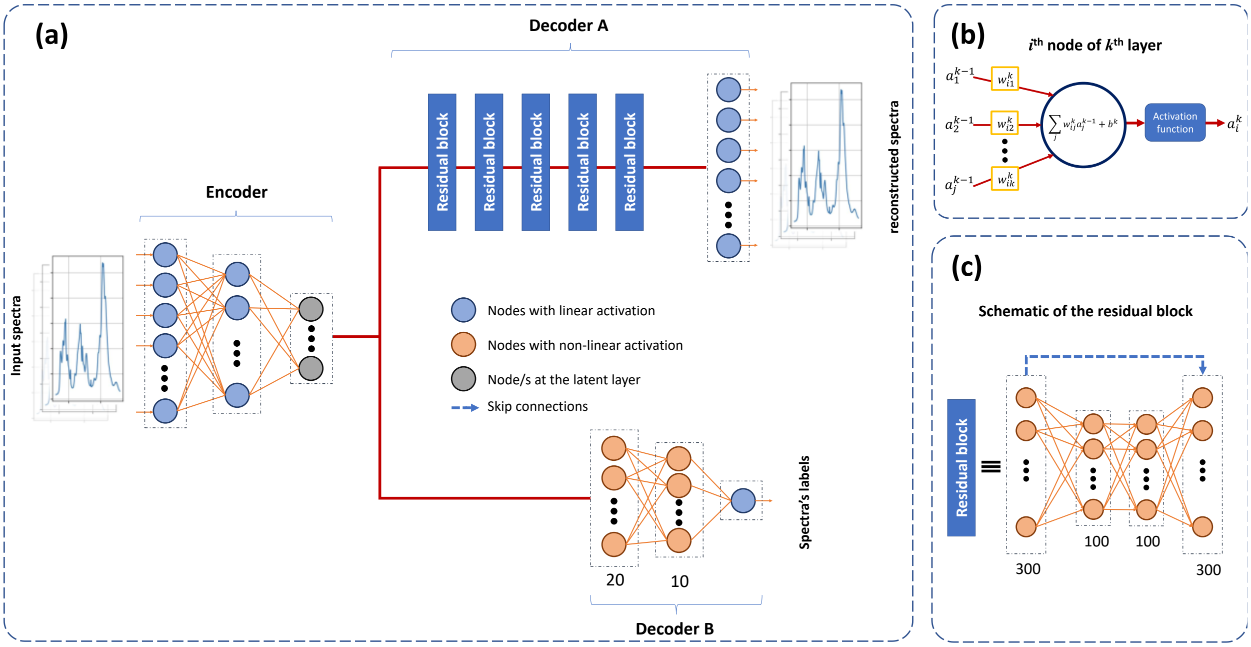


**Supplementary Figure 1.** (a) Schematic of the proposed network architecture consisting of one linear encoder and two parallel decoders. Decoder A expands the latent layer dimension using a stack of residual blocks to the dimension of the input and tries to reconstruct the input, while decoder B matches the latent values to the data's labels. (b) Connections and topology of each node in all dense layers and (c) the architecture of the residual block used in the decoder A.

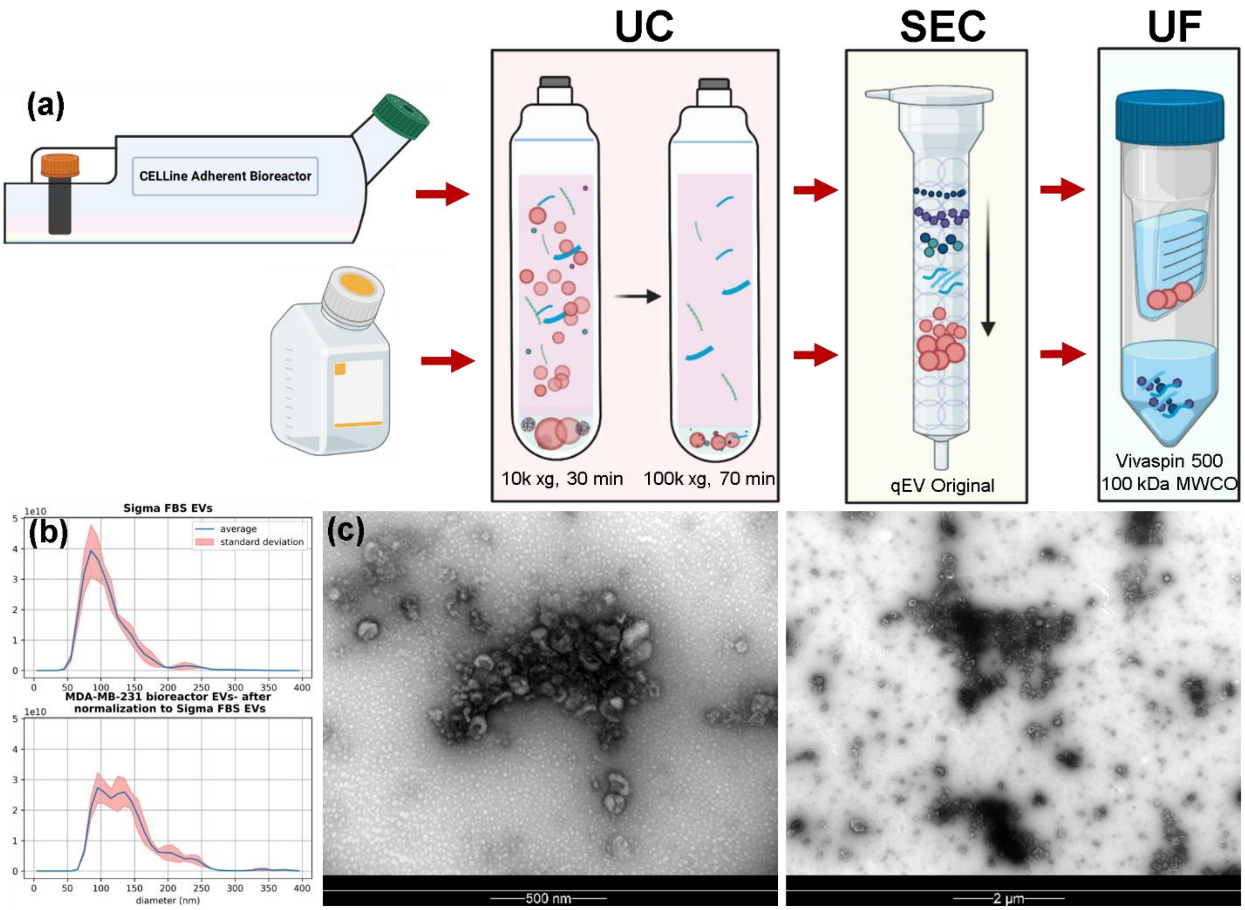


**Supplementary Figure 2.** EV mixture model used for SERS detection showing (a) EVs from an MDA-MB-231 bioreactor culture and commercial FBS isolated using differential ultracentrifugation (UC) followed by size exclusion chromatography (SEC) and ultrafiltration (UF), (b) NTA data from both samples after buffer exchange and normalizing particle concentration for SERS experiments, and (c) TEM images of bioreactor-produced MDA-MB-231 small EVs at different magnifications.

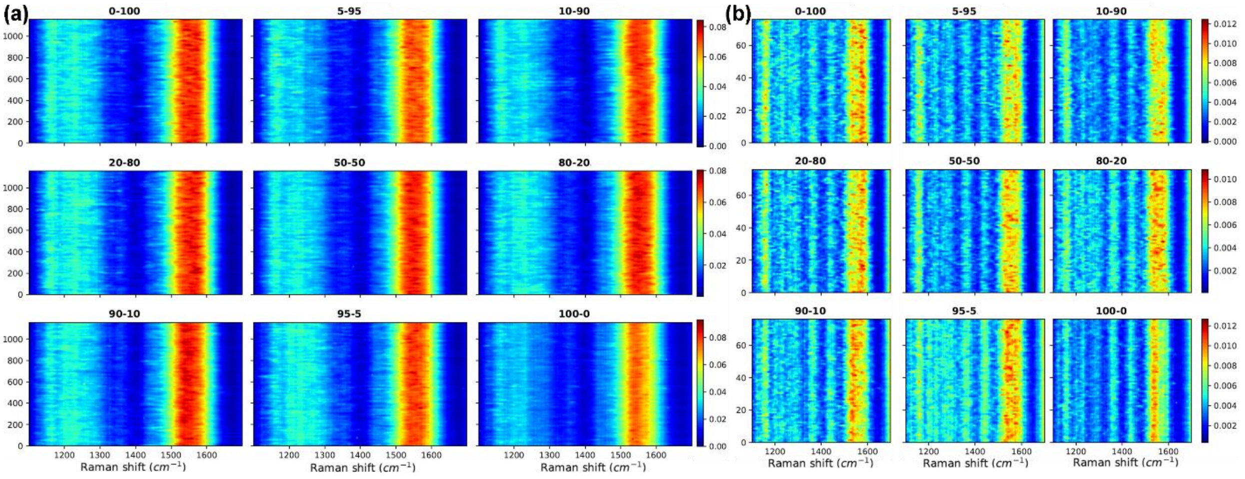


**Supplementary Figure 3.** Preprocessed Raman spectra from EV spike-in mixture model showing (a) all the acquired Raman spectra and (b) the same spectra following averaging, and excessive baseline removals for better illustrating the heterogeneity in each sample.

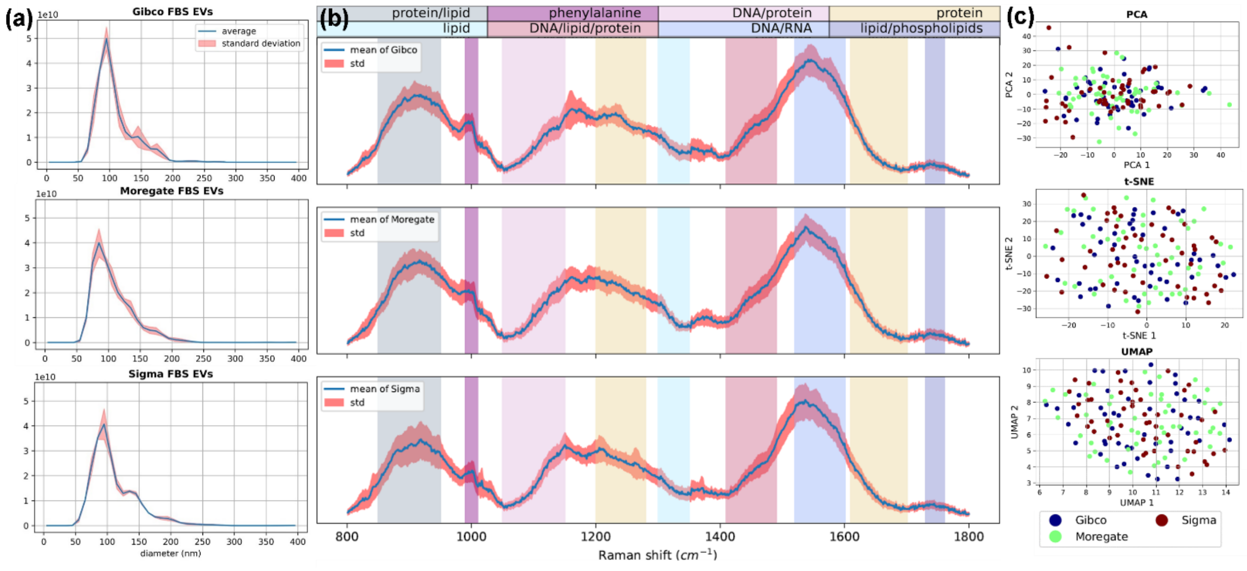


**Supplementary Figure 4.** Comparison of Gibco, Moregate, and Sigma FBS-derived small EVs using SERS showing (a) NTA data after normalizing particle concentration following buffer exchange, (b) Raman spectra with color-coded general peak associations, and (c) PCA, t-SNE, and UMAP dimensionality reduction of FBS EV Raman spectra showing no apparent clustering.

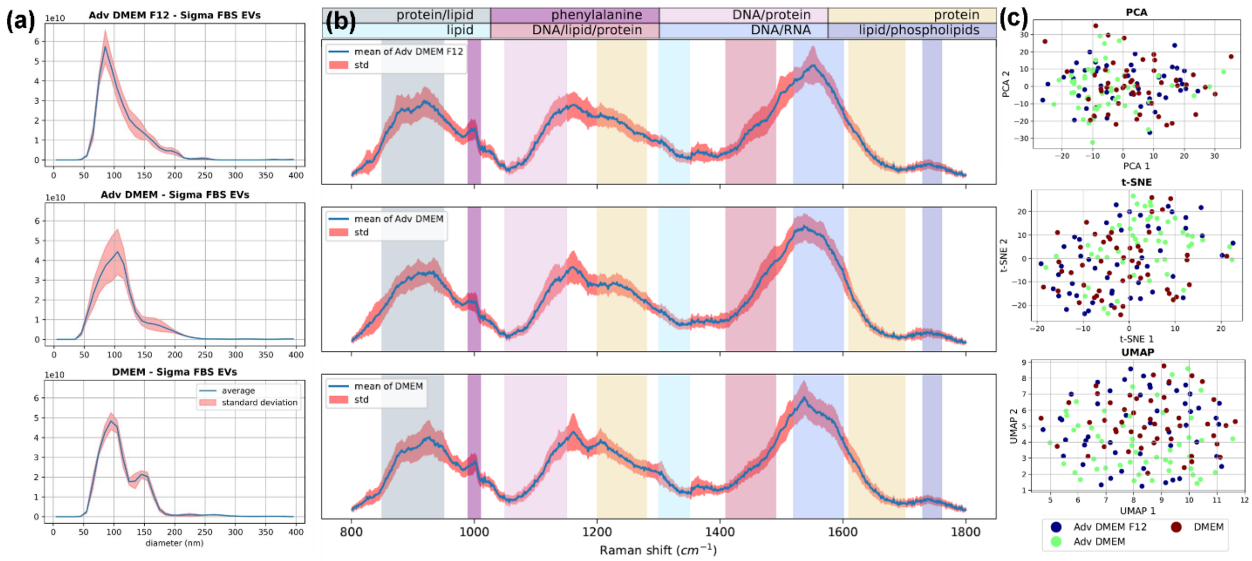


**Supplementary Figure 5.** Comparison of Sigma FBS-derived small EVs stored in Advanced DMEM/F12, Advanced DMEM, or DMEM for 2 weeks at 4 ˚C using SERS showing (a) NTA data after normalizing particle concentration following buffer exchange, (b) Raman spectra with color-coded general peak associations, and (c) PCA, t-SNE, and UMAP dimensionality reduction of spectra for Sigma FBS-derived small EVs showing no apparent clustering.
